# Demographic inference using a particle filter for continuous Markov Jump processes

**DOI:** 10.1101/382218

**Authors:** Donna Henderson, Sha (Joe) Zhu, Chris Cole, Gerton Lunter

**Affiliations:** Wellcome Centre for Human Genetics, Roosevelt Drive, Oxford OX3 7BN, UK; Big Data Institute, Old Road, Oxford OX3 7FZ, UK; MRC Weatherall Institute of Molecular Medicine, John Radcliffe Hospital, Headington, Oxford Oxford OX3 9DS, UK

**Keywords:** Continuous-time continuous-space Markov jump process, Particle filter, Sequential Monte Carlo, Variational Bayes, Demo-graphic Inference

## Abstract

Demographic events shape a population’s genetic diversity, a process described by the coalescent-with-recombination (CwR) model that relates demography and genetics by an unobserved sequence of genealogies. The space of genealogies over genomes is large and complex, making inference under this model challenging.

We approximate the CwR with a continuous-time and -space Markov jump process. We develop a particle filter for such processes, using way-points to reduce the problem to the discrete-time case, and generalising the Auxiliary Particle Filter for discrete-time models. We use Variational Bayes for parameter inference to model the uncertainty in parameter estimates for rare events, avoiding biases seen with Expectation Maximization.

Using real and simulated genomes, we show that past population sizes can be accurately inferred over a larger range of epochs than was previously possible, opening the possibility of jointly analyzing multiple genomes under complex demographic models.

Code is available at https://github.com/luntergroup/smcsmc

**MSC 2010 subject classifications:** Primary 60G55, 62M05, 62M20, 62F15; secondary 92D25.

## 1. Introduction

The demographic history of a species has a profound impact on its genetic diversity. Changes in population size, migration and admixture events, and population splits and mergers, shape the genealogies describing how individuals in a population are related, which shape the pattern and frequency of observed genetic polymorphisms in extant genomes. By modeling this process and integrating out the unobserved genealogies, it is possible to infer the population’s demographic history from the observed polymorphisms. However, in practice this is challenging, as individual mutations provide limited information about tree topologies and branch lengths. If many mutations were available to infer these genealogies this would not be problematic, but the expected number of observed mutations increases only logarithmically with the number of genomes, and recombination causes genealogies to change along the genome at a rate proportional to the mutation rate. As a result there is considerable uncertainty about the genealogies underlying a sample of genomes, and because the space of genealogies across the genome is vast, integrating out this latent variable is hard.

A number of approaches have been proposed to tackle this problem [reviewed in 52]. A common approximation is to treat recombination events as known and assume unlinked loci, either by treating each mutation as independent [2, 48, 3, 26, 25, 16], or by first identifying tracts of genetic material unbroken by recombination [15, 6, 46, 28, 29]. To account for recombination while retaining power to infer earlier demographic events, it is necessary to model the genealogy directly. ARGWeaver [49] implements the full coalescent with recombination (CwR) model and uses Markov chain Monte Carlo (MCMC) for inference. While this works well, ARGWeaver does not allow the use of a complex demographic model, and since mutations are only weakly informative about genealogies this leaves the inferred trees biased towards the prior model and less directly suitable for inferring demography. Restricting to single diploid genomes, the Pairwise Sequentially Markovian Coalescent (PSMC) model [35] uses an elegant and efficient inference method, but with limited power to detect recent changes in population size or complex demographic events. Several other approaches exist that improve on PSMC in various ways [53, 51, 57, 60], but remain limited particularly in their ability to infer migration.

A promising approach which so far has not been applied to this problem is to use a particle filter. Particle filters have many desireable properties [23, 12, 1, 14], and applications to a range of problems in computational biology have started to appear [59, 55, 21, 61]. Like MCMC methods, particle filters converge to the exact solution in the limit of infinite computational resources, are computationally efficient by focusing on realisations that are supported by the data, do not require the underlying model to be approximated, and generate explicit samples from the posterior distribution of the latent variable. Unlike MCMC, particle filters do not operate on complete realisations of the model, but construct samples sequentially, which is helpful since full genealogies over genomes are cumbersome to deal with.

Originally, particle filters were introduced for models with discrete time evolution and with either discrete or continuous state variables [50, 23]. Here, the latent variable is a sequence of piecewise constant genealogical trees along the genome, with trees changing only after recombination events that, in mammals, occur once every several hundred nucleotides. The observations of the model are polymorphisms, which are similarly sparse. Realizations of the discrete-time model of this process (with “time” playing the role of genome position) are therefore stationary and silent at most transitions, leading to inefficient algorithms. Instead, it seems natural to model the system as a Markov jump process (or purely discontinuous Markov process, [18]), a continuous-time stochastic process with as realisations piecewise constant functions *x* : [1, *L*] ↦𝕋, where 𝕋 is the state space of the Markov process (here the space of all genealogical trees over a given number of genomes) and *L* the length over which observations are made (here the genome size).

Particle filters have been generalised to continuous-time diffusions [10, 22, 17], as well as to Markov jump processes on discrete state spaces [44, 42], and hybrids of the two [13, 54], as well as to piecewise deterministic processes [63]; for a general treatment see [11, 9]. Here we focus on the case of Markov jump processes that are continuous in both time and space. The algorithm we propose relies on Radon-Nikodym derivatives [see e.g. 17], and we establish criteria for choosing a finite set of “waypoints” that makes it possible to reduce the problem to the discrete-time case, while ensuring that particle depletion remains under control.

Although the algorithm generally works well, we found that for the CwR model we obtain biased inferences for some parameters whose associated latent variables have long forgetting times, requiring fixed-lag smoothing with long lags resulting in particle degeneracy [45]. For discrete-time models the Auxiliary Particle Filter [47] addresses a related problem by “guiding” the sampler towards states that are likely to be relevant in future iterations, using an approximate likelihood that depends on data one step ahead. This approach does not work well for some continuous-time models, including ours, that have no single preferred time scale. Instead we introduce an algorithm that shapes the resampling process by an approximate “lookahead likelihood” that can depend on data at arbitrary distances. Using simulations we show that this substantially reduces the bias.

The particle filter generates samples from the posterior distribution of the latent variable, and we infer the model parameters from this sample. One strategy is to use stochastic expectation-maximization [SEM; 43]. An issue with this approach is that the joint posterior distribution over parameters is represented by a point-mass, ignoring uncertainty in parameter estimates. Combined with the bias due to self-normalized importance sampling which cause particle filters to under-sample low-rate events, this result in a non-zero probability of inferring a zero event rate, which is a fixed point of an SEM procedure. In principle this could be avoided by using an appropriate prior on the rate parameters. To implement this we use Variational Bayes to estimate an approximate joint posterior distribution over parameters and latent variables, partially accounting for the uncertainty in the inferred parameters, as well as providing way to explicitly include a prior. In this way zero-rate estimates are avoided, and more generally we show that this approach further reduces the bias in parameter estimates.

Applying these ideas to the CwR model, we find that the combination of lookahead filter and Variational Bayes inference enables us to analyze four diploid human genomes simultaneously, and infer demographic parameters across epochs spanning more than 3 orders of magnitude, without making model approximations beyond passing to a continuous-locus model.

The remainder of the paper is structured as follows. We first introduce the particle filter, generalise it to continuous-time and -space Markov jump processes, describe how to choose waypoints, introduce the lookahead filter, and describe the Variational Bayes procedure for parameter inference. In the results section we first introduce the continuous-locus CwR process, then discuss the lookahead likelihood, choice of waypoints and parameter inference for this model, before applying the model to simulated data, and finally show the results of analyzing sets of four diploid genomes of individuals from three human populations. A discussion concludes the paper.

## 2. Methods

### 2.1. Particle filters

Particle filters methods, also known as Sequential Monte Carlo (SMC) [14], generate samples from complex probability distributions with high-dimensional latent variables. An SMC method uses importance sampling (IS) to approximate a target distribution using weighted random samples (particles) drawn from a tractable distribution. We briefly review the discrete-time case. Suppose that particles {(*x*^(*i*)^, *w*^(*i*)^)} _*i*=1,*…,N*_, approximate a distribution with density *p*(*x*), such that

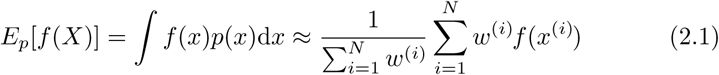

for any bounded continuous function *f*, where *X* ∼ *p*(*x*)d*x*. We use “approximate” and ≈ to mean that 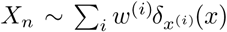 converges in distribution to *X* ∼ *p*(*x*)d*x*, so that equality holds in (2.1) as *N* → *∞*. Under some conditions, we can use IS to obtain particles approximating another distribution *q*(*x*)d*x*:

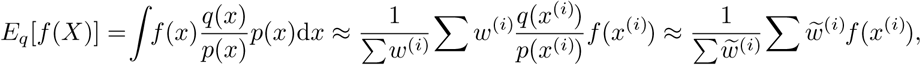

Where 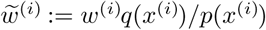, and the last step holds because 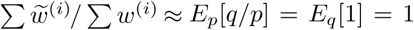. This shows that 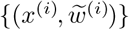 approximate *q*(*x*)d*x*. Note that any constant factor in *w*^(*i*)^ drops out because of the normalisation, so that it is sufficient to determine the ratio *q*(*x*)*/p*(*x*) up to a constant. A particle filter builds the desired distribution sequentially, making it suited to hidden Markov models, for which the joint distribution of latent variables *X* and observations *Y* (both taken to be discrete in this section) have the form

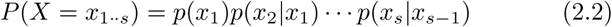

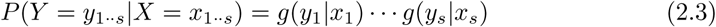

Here 1*··s* denotes the set {1, 2, …, *s*}, and *x* = *x*_1*··s*_ = (*x*_1_, *x*_2_, …, *x*_*s*_) and *y* are vectors. Let {(*x*^(*i*)^, *w*^(*i*)^)} be particles approximating the posterior *P* (*X*_1*··s*_ = *x*_1*··s*_|*Y*_1*··s*_ = *y*_1*··s*_), which for brevity we write as *P* (*x*_1*··s*_|*y*_1*··s*_). If 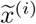is the vector obtained by extending *x*^(*i*)^ with a sample from *P* (*x*_*s*+1_|*x*^(*i*)^), then from (2.2) and (2.3) it follows that 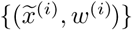 approximate *P* (*x*_1*··s*+1_|*y*_1*··s*_) ∝ *P* (*x*_1*··s*+1_, *y*_1*··s*_). Now, *P* (*x*_1*··s*+1_|*y*_1*··s*+1_) ∝ *P*(*x*_1*··s*+1_, *y*_1*··s*+1_) = *P*(*x*_1*··s*+1_, *y*_1*··s*_)*g*(*y*_*s*+1_|*x*_*s*+1_), so that using IS and setting

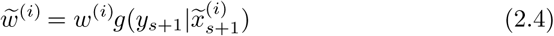

we obtain particles 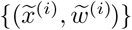 that approximate *p*(*x*_1*··s*+1_|*y*_1*··s*+1_). This shows how to sequentially construct particles that approximate the posterior distribution *p*(*x*_1*··L*_|*y*_1*··L*_). Instead of sampling from *p*(*x*_*s*+1_|*x*^(*i*)^), any proposal distribution *q*(*x*_*s*+1_|*x*^(*i*)^, *y*_1*··L*_) (subject to conditions) can be used, which is advantageous if *q* is easier to sample from, is closer to the posterior distribution, or has heavier tails than *p*. Again, IS accounts for the change in sampling distribution, resulting in

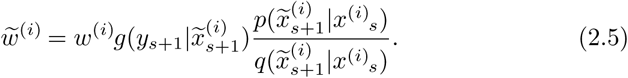

For now we will choose *q* to be independent of *y*. Because samples from *q* do not follow the desired posterior *p*(*x*|*y*), the fraction of particles close to the posterior’s mode diminishes exponentially at each iteration until (2.1) fails altogether. To address this, samples are drawn from the approximating distribution itself, assigning each resampled particle weight 1*/N* – interestingly, this is the same process that is used in the Wright-Fisher model with selection to describe how fitness differences shape an evolving population [30]. This procedure tends to remove particles that have drifted from the mode of the posterior and have low weight, and duplicates particles with large weights, while preserving the validity of (2.1). Although resampling substantially decreases the future variance of (2.1), it increases the variance at the present iteration. To avoid increasing this variance unnecessarily, resampling is performed only when the estimated sample size *ESS* = (Σ *w*^(*i*)^)^2^*/* (*w*^(*i*)^)^2^ drops below a threshold, often *N/*2. In addition, we use systematic resampling to minimize the variance introduced when resampling is performed [7]. This leads to Algorithm 1 [23].

The algorithm generates an approximation to *p*(*x*_1*··s*_|*y*_1*··s*_) rather than *p*(*x*_*s*_|*y*_1*··s*_), but we follow [14] in calling it a particle filter algorithm instead of a smoothing algorithm (although our use of fixed-lag distributions for parameter estimation, Section 3.4, is a partial smoothing operation). The marginal likelihood can be estimated (although with high variance, see [4]) by setting the weights to 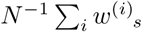 rather than *N* ^*−*1^ when particles are resampled. This makes the weights asymptotically normalized, so that (2.1) becomes 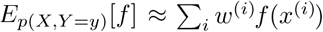, and 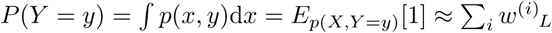.

#### Algorithm 1 Particle filter

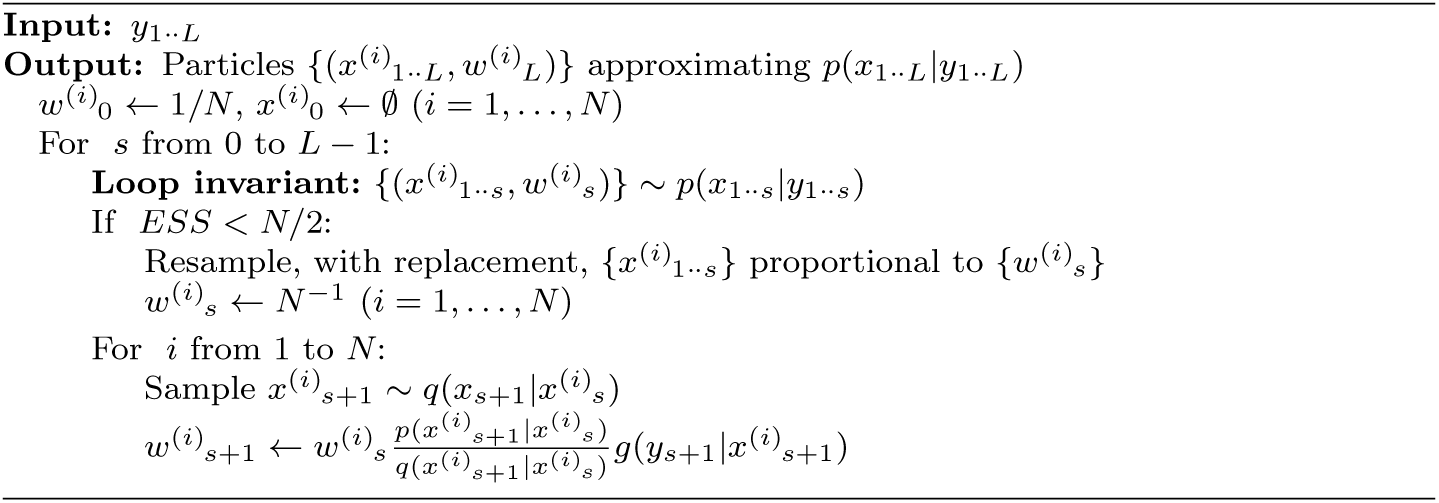

### 2.2. Continuous-time and -space Markov jump processes

We now consider Markov jump processes that have as realisations piecewise constant functions *x* : [1, *L*) ↦ 𝕋 where 𝕋 is the state space of the Markov process. We consider the case that 𝕋 is uncountable, e.g. 𝕋 = ℝ^*n*^ (our main interest is in the case 𝕋 = ℝ^*n−*1^ *× T*_*n*_ of genealogical trees over *n* sequences; *T*_*n*_ is the set of rooted tree topologies over *n* tips). Let (*𝒳, ℱ*_*x*_, *π*_*x*_) be a probability space with *𝒳* = 𝕋^[1,*L*)^ the space of possible realisations of the stochastic process *X* = {*X*_*s*_} _*s*∈[1,*L*)_, *ℱ*_*x*_ *⊂ 𝒫* (*𝒳*) the *σ*-algebra of events, and *π*_*x*_(*X*) the probability measure on induced by the stochastic process *X*. The definition of a Markov model for uncountable phase space 𝕋 involves the concept of conditional distributions; see the Appendix for details. For the hidden Markov model, assume *X* is not observed, and that observations *Y* are generated by a point process whose intensity at time *s* depends on *X*_*s*_ [a Cox process, see e.g. 34]. We define the space of observations *Y* = *𝒫*_*<*ℕ_([1, *L*) *× M*) of finite subsets of [1, *L*) *× M*, where *M* is a discrete set of potential events that may occur at some *s ∈* [1, *L*). For an element 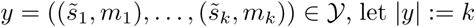 be the number of events in the full observation, and let *λ*(*y*) be the Lebesgue measure (d*s*)^|*y*|^, so that the “emission” distribution *π*(*Y* |*X* = *x*) has a density *h*(*y*|*x*) relative to *λ*; for Cox processes this density factorizes as 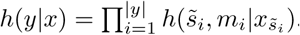. The probability space for the joint process is (*𝒳 × 𝒴, ℱ, π*), and the posterior distribution of interest is *π* conditioned on an observation *y* ∈ 𝒴, written as *π*(*X*|*Y* = *y*).

To describe the Markov jump process version of algorithm 1 we introduce some more notation. As above *π*_*x*_ denotes the prior distribution of the latent variable *X*, and *ξ*_*x*_ denotes the proposal distribution, both Markov processes on *X*, playing the role of *p*(*x*) and *q*(*x*) for the discrete case. Let *a*:*b* denote the real interval [*a, b*), and let *α*^*a*:*b*^ denote the restriction of a measure or function *α* to the interval [*a, b*) ⊆ [1, *L*); similarly *y*_*a*:*b*_ := *y ∩* ([*a, b*) × *M*) and *X*_*a*:*b*_ := {*X*_*s*_} _*s*∈[*a,b*)_. The particle filter algorithm uses the notation (d*α/*d*β*)(*x*) for distributions *α* and *β* to denote their Radon-Nikodym derivative: the ratio of their density functions with respect to a common reference measure, evaluated at *x*. To simplify notation we write the Radon-Nikodym derivative of two conditional distributions *α*(*X*| *𝒢*) and *β*(*X*| *𝒢*) at *x* as (d*α/*d*β*)(*x*| *𝒢*), and we also do not explicitly restrict distributions to their appropriate intervals when this is clear from the context, so that we write for example 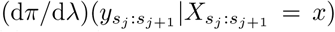 instead of 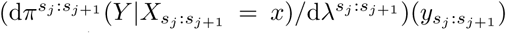. With this notation we can formulate Algorithm 2.

#### Algorithm 2 Particle filter for Markov jump processes

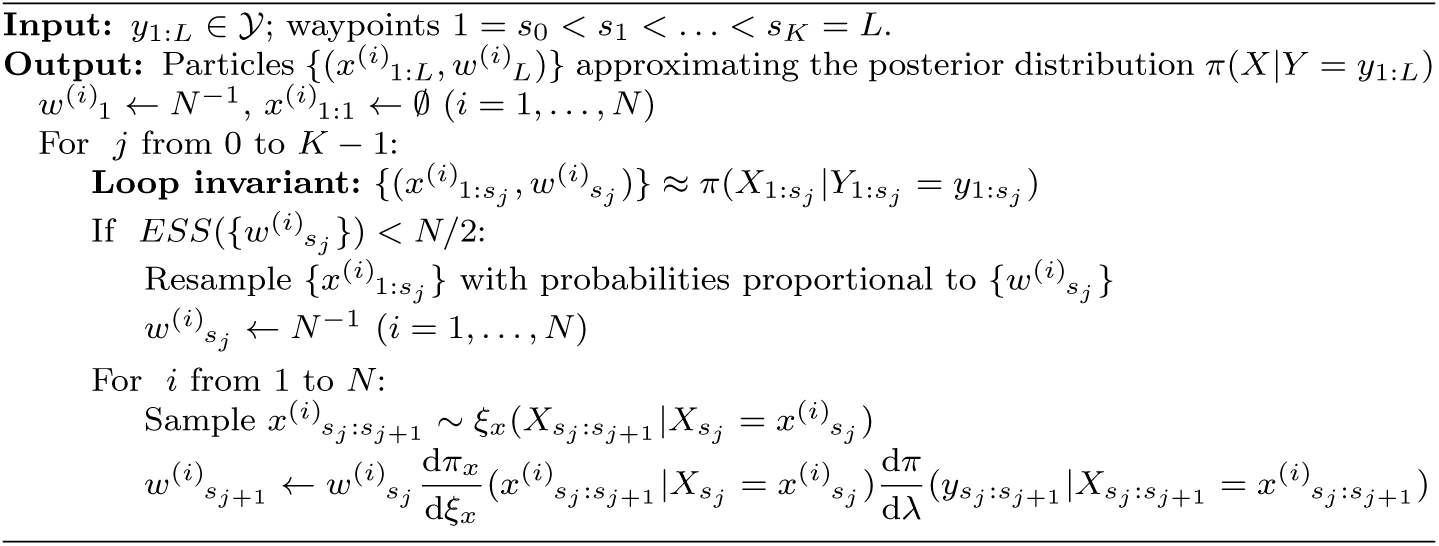

The choice of waypoints *s*_1_, …, *s*_*K*_ is discussed in section 2.3 below; in particular they need not coincide with the event times 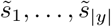 in the observation *y*. Note that there is no initialization step; instead, initially *x*^(*i*)^_1.1_ = ∅, and the first sample will be drawn from *ξ* conditioned on an empty set, i.e. the unconditional distribution. The loop invariant is trivially true when *j* = 0 since 1:*s*_0_ = ∅. As with Algorithm 1 it is possible to estimate the likelihood density *π*_*θ*_(*y*_1:*L*_) by replacing the factors *N* ^*−*1^ with 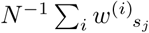; then the likelihood density w.r.t. *λ*(d*y*) = (d*s*)^|*y*|^ is approximated by Σ_*i*_ *w*^(*i*)^_*L*_.

Note that by the nature of Markov jump processes, particles that start from identical states have a positive probability of remaining identical after a finite time. Combined with resampling, this causes a considerable number of particles to have one or more identical siblings. For computational efficiency we represent such particles once, and keep track of their multiplicity *k*. When evolving a particle with multiplicity *k >* 1, we increase the exit rate *k*-fold, and when an event occurs one particle is spawned off while the remaining *k −*1 continue unchanged.

The absence of events along a time interval is also an “observation” and is assigned a likelihood density. In practice some intervals may not be observable for technical reasons. We assume that the observability process is independent of the Markov jump process, so that the likelihood density d*π/*d*λ* can be set to 1 across these intervals.

### 2.3. Choosing waypoints

The choice of waypoints *s*_*j*_ can significantly impact the performance of the algorithm: choosing too few increases the variance of the approximation, and choosing too many slows down the algorithm without increasing its accuracy. Waypoints affect where the algorithms consider whether or not to resample. A high density of waypoints is therefore always acceptable, but a low density may result in particle degeneration. Choosing a waypoint at every event ensures that any weight variance induced at these sites is mitigated, but there is still the opportunity for weight variance to build up between events. Here we derive a criterion on the waypoints that limits this effect.

Let *R*(*s*) = *f* (*X*_*s*_) be the stochastic variable that measures the instantaneous rate of occurrence of events for a particular (random) particle *X*, and let 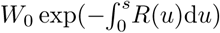 be that particle’s time-dependent weight; the dependence on *X* is not written explicitly. Note that the expression for *W* (*s*) is valid as long as no events have occurred in the interval [0, *L*). We assume that *R*(*s*) is time-homogeneous, that it can be approximated by a Gaussian process, that particles are drawn from the equilibrium distribution, and that *W*_0_ and *R*(*s*) are independent. Write ⟨*V* (*X*) ⟩ := ∫*V* (*X*)d*π*(*X*) for the expectation of *V* over *π*(*X*). Writing *R*(*s*) = *µ* + *r*(*s*) where *µ* = ⟨*R*(*s*)⟩ is the mean event rate (which is independent of *s* by assumption), then

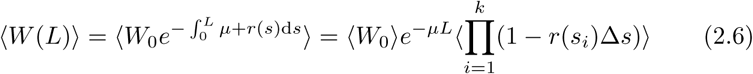

as *k* → *∞*, where Δ*s* = *L/k* and *s*_*i*_ = *i*Δ*s*. The last expectation becomes

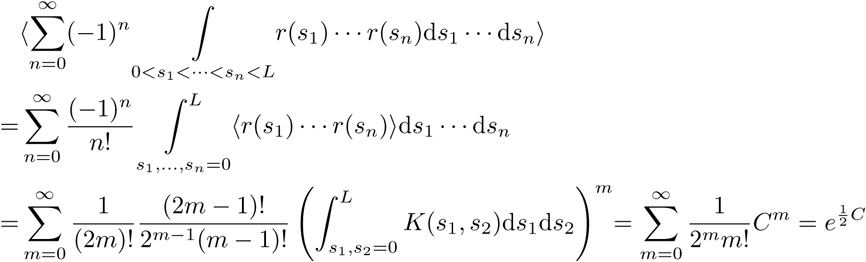

where in the second equality we used the formula for higher moments of a Gaussian distribution, *K* is the covariance function of the Gaussian process *r*(*t*), and *C* is the integral 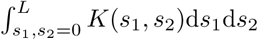. Now define *σ*^2^ := *K*(*s, s*) and assume that the covariance function satisfies 0 *≤ K*(*s*_1_, *s*_2_) *≤ σ*^2^, then 0 *≤ C ≤ σ*^2^*L*^2^ and

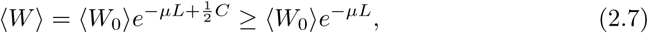

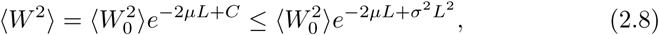

so that across an interval [0, *L*) where no events occur,

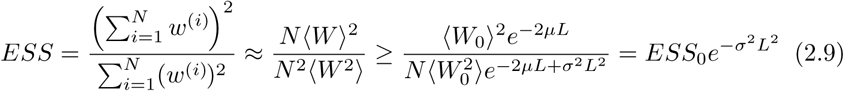

where *ESS*_0_ is the expected sample size at *s* = 0, and ≈ denotes convergence in distribution as *N* → *∞* as before. Therefore, if we choose waypoints at every event, adding additional waypoints so that they are never more than a distance 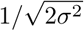 apart, the ESS will not drop more than a factor 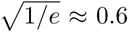 between waypoints, and particle degeneracy is avoided.

In practice particles will not be drawn from the equilibrium distribution *π*_*x*_(*X*), but from the joint distribution on *X* and *Y* conditioned on observations *y*. However, for most likelihoods conditioning will reduce the variance of *R* as observations tend to constrain the distribution of likely particles, making this a conservative assumption. The other assumption that is likely not met is that *R*(*t*) is a Gaussian process; it is less clear whether making this approximation will in practice be conservative. Therefore the minimum waypoint density derived here serves as a guide; breakdown of the assumptions made here can be monitored by tracking the *ESS*, increasing the density of waypoints if necessary.

### 2.4. A lookahead particle filter for Markov jump processes

At the *j*th iteration, algorithm 2 uses data up to waypoint *s*_*j*_ to build particles approximating 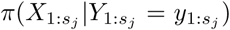. This is reasonable as 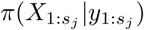 is independent of data beyond *s*_*j*_. However, not all particles are equally important for approximating subsequent posteriors. Emphasising particles that will be relevant in future at the expense of those relevant only to 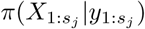 would improve the overall accuracy of the algorithm. This argument echoes the justification of resampling: although resampling increases the variance of the approximation to the current partial posterior, it reduces the variance at subsequent iterations by increasing the number of particles that are likely to contribute to future distributions. The Auxiliary Particle Filter [47] implements this intuition for discrete-time models *p*(*X*_1:*n*_, *Y*_1:*n*_) by targeting a special resampling distribution [33]. This distribution includes a “lookahead” factor 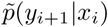 that approximates the probability of observing data *y*_*i*+1_ at the next time step given the current state *x*_*i*_. Importance sampling is used to keep track of the desired distribution *p*(*X*_1:*i*_ | *Y*_1:*i*_).

In the continuous-time context it is natural to look an arbitrary distance ahead, instead of using a one-step or fixed-distance lookahead. Since at intermediate stages of the algorithm no approximation of the future state is yet available, the lookahead distribution can be conditioned on the current state only, and will in practice be an approximation of the true distribution. It should be heavy-tailed with respect to the true distribution to ensure that the variance of the estimator remains finite [14], which implies that the distribution should not depend on data too far beyond *s*; what is “too far” depends on how well the lookahead distribution approximates the true distribution.

The lookahead distribution is only evaluated on a fixed observation *y*, and is not used to define a distribution over *y*, but to quantify the plausibility of a current state *x*^(*i*)^ _s_. For this reason we refer to it as the lookahead *likelihood*. In fact, for the validity of the algorithm below it is not necessary for this likelihood to derive from a proper distribution. We therefore define the lookahead likelihood as a family of functions *h*^*s*^(*y*_*s*:*L*_|*x*_*s*_) : *Y*_*s*:*L*_ *×* 𝕋 → ℝ, and an associated family of unnormalized distributions 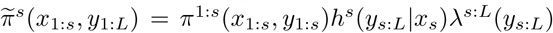 on. The functions *h*^*s*^ can be chosen arbitrarily except that *h*^*s*^ (·, *x*_s_)*λ*_*s*:*L*_ must be absolutely continuous w.r.t. *π*^*s*:*L*^(*·*|*X*_*s*_ = *x*_*s*_) to ensure that importance sampling is justified.

The lookahead algorithm 3 keeps track of two sets of weights and one set of samples, together forming two sets of particles that approximate the resampling and target distributions. The resampling distribution alone controls the resampling process, and ensures that samples that are likely to contribute to subsequent posteriors, as quantified by the lookahead likelihood, are likely to be kept even if they have low likelihood under the current target distribution.

#### Algorithm 3 Markov-jump particle filter with lookahead

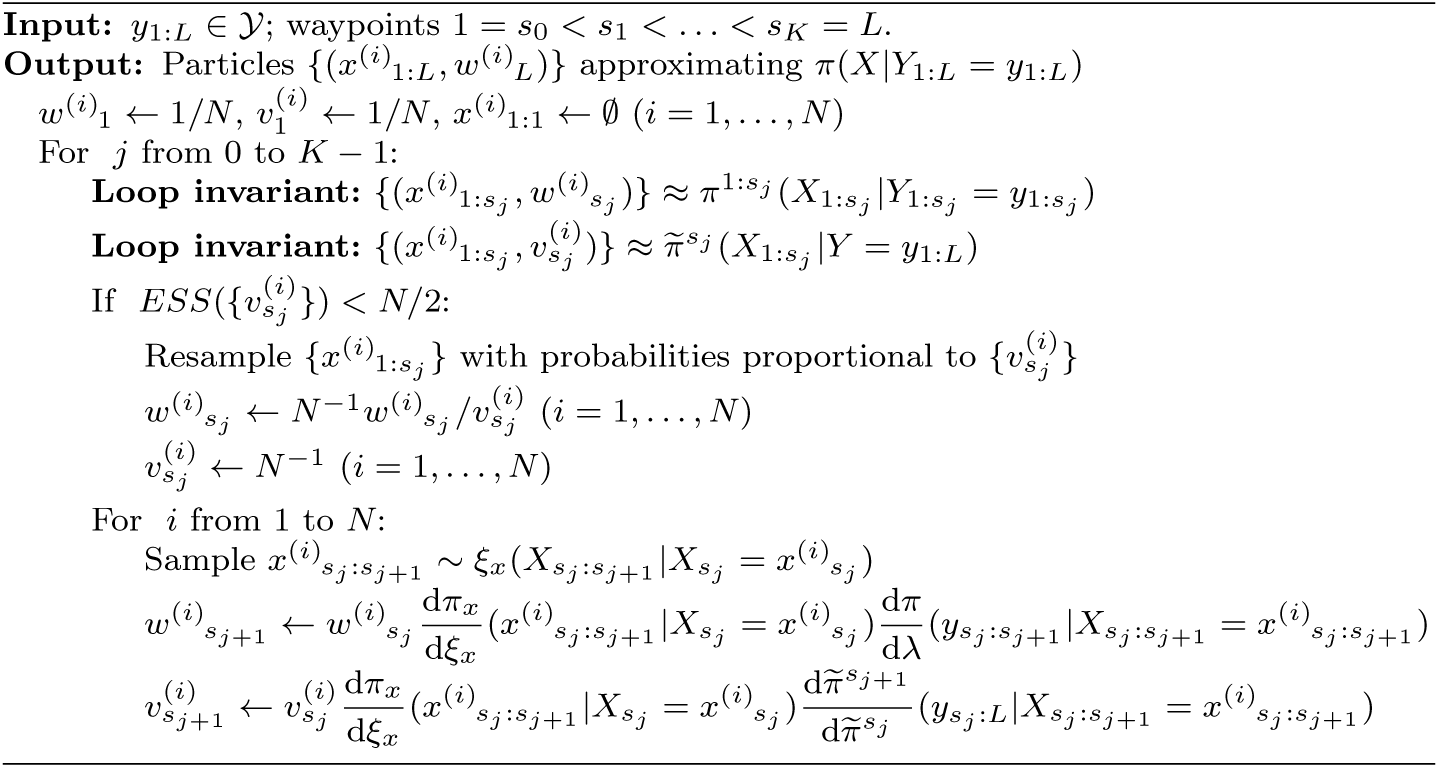

### 2.5. Parameter inference

Parameters can be inferred by stochastic expectation maximization (SEM), which involves maximizing the expected log likelihood over the posterior distribution of latent variables. Note that the expected log likelihood for a Poisson process with rate *θ* is *c* log *θ − qθ* where *c* and *q* are the expected event count and opportunity, which is maximized for *θ* = *c/q*. We consider Markov jump processes *X*_*s*_ parameterized by a vector of parameters *θ*, for which the distribution *π*_*x*_(d*x*|*θ*) can be written as

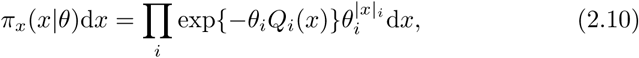

where |*x*_*i*_| is the event count and *Q*_*i*_(*x*) is the total opportunity, for events of type *i* in realisation *x*; both can be random variables. Similar to the Poisson case, the parameters maximizing the expected log likelihood are

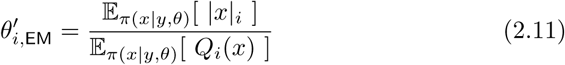

The expectations can be computed by using samples over *x* ∼ *π*(*x* |*y, θ*) as approximated by algorithm 3.

Particularly in cases where some event types are rare, Variational Bayes can improve on EM by iteratively estimating posterior distributions rather than point estimates of *θ*. A tractable algorithm is obtained if the joint posterior *π*(*x, θ*|*y*)d*x*d*θ* is approximated as a product of two independent distributions over *x* and *θ*, and an appropriate prior over *θ* is chosen. For the Poisson example above, combining a Γ(*θ*|*α*_0_, *β*_0_) prior with the likelihood *θ*^*c*^*e*^*−qθ*^ results in a Γ(*θ*|*α*_0_ + *c, β*_0_ + *q*) posterior. Similarly, with this choice the Variational Bayes approximation results in an inferred posterior distribution of the form

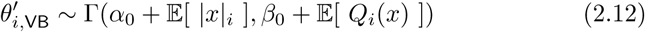

where expectations are taken over *x* ∼ *∫π*(*x*|*y, θ*)*π*(*θ*)d*θ*, and *π*(*θ*) is the current posterior over *θ*. It would appear that obtaining samples from this distribution is intractable. However, if *π*(*θ*) is a Gamma distribution, *θ* can be integrated out analytically in the likelihood *π*(*x, y*|*θ*)Γ(*θ*|*α, β*), resulting in an expression that is identical to the likelihood of the point estimate *θ*_*i*_ = *α*_*i*_*/β*_*i*_ except for an additional scaling factor 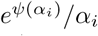 for each event of type *i* in *x*; here *Ψ* is the digamma function, see the Appendix for details. These scaling factors render the normalization constant of the likelihood intractable, but fortunately SMC algorithms only require likelihoods up to normalization so that algorithm 3 can be used to generate samples from this distribution at no additional computational cost.

## 3. Results

### 3.1. The sequential coalescent with recombination model

The coalescent-with-recombination (CwR) process, and the graph structure that results from it, was first described by [32], and was given an elegant mathematical description by [24], who named the resulting structure the Ancestral Recombination Graph (ARG). Like the coalescent process, these models run backwards in time and consider the entire sequence at once, making it difficult to use them for inference on whole genomes. The first model of the CwR process that evolves spatially rather than in the time direction was introduced by [64], opening up the possibility of inference over very long sequences. Like Griffiths’ process, the Wiuf-Hein algorithm operates on an ARG-like graph, but it is more efficient as it does not include many of the non-observable recombination events included in Griffiths’ process. The Sequential Coalescent with Recombination Model (SCRM) developed by [56] further improved efficiency by modifying Wiuf and Hein’s algorithm to operate on a local genealogy rather than an ARG-like structure. Besides the “local” tree over the observed samples, this genealogy includes branches to non-contemporaneous tips that correspond to recombination events encountered earlier in the sequence. Recombinations on these “non-local” branches can be postponed until they affect observed sequences, and can sometimes be ignored altogether, leading to further efficiency gains while the resulting sample still follows the exact CwR process. An even more efficient but approximate algorithm is obtained by culling some non-local branches. In the extreme case of culling *all* non-local branches the SCRM approximation is equivalent to the SMC’ model [40, 38]. With a suitable definition of “current state” (i.e., the local tree including all non-local branches) these are all Markov processes, and can all be used in the Markov jump particle filter; here we use the SCRM model with tunable accuracy as implemented in [56].

The state space 𝕋 of the Markov process is the set of all possible genealogies at a given locus. The probability measure of a realisation *x* can be written as *π*_*x*_(*x*|*ρ, C*)d*x* =

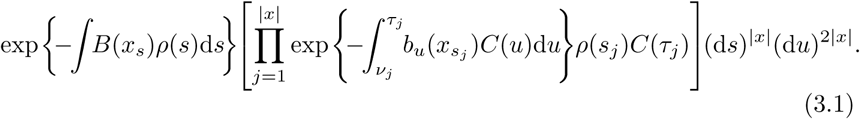

Here *x* ∈ *X* is the sequence of genealogies; |*x*| is the number of recombinations that occurred on *x*; *b*_*u*_(*x*_*s*_) is the number of branches in the genealogy at position *s* at time 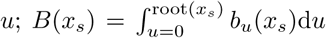is the total branch length of *x*_s_ ; *ρ*(*s*) is the recombination rate per nucleotide and per generation at locus *s*, so that *ρ*(*s*)*B*(*x*_*s*_) is the exit rate of the Markov process in state *x*_*s*_; (*s*_*j*_, *v*_*j*_) is the locus and time of the *j*th recombination event; *τ*_*j*_ *> v*_*j*_ is the time of the corresponding coalescence event; and *C*(*u*) = 1*/*2*N*_*e*_(*u*) is the coalescence rate in generation *u*. See Section 5.3 for more details.

Mutations follow a Poisson process whose rate at *s* depends on the latent variable *x*_*s*_ via *µ*(*s*)*B*(*x*_*s*_) where *µ*(*s*) is the mutation rate at *s* per nucleotide and per generation. Mutations are not observed directly, but their descendants are; a complete observation is represented by a set 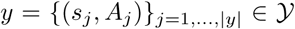 where *s*_*j*_ ∈ [1, *L*) is the locus of mutation *j*, and *A*_*j*_ ∈{0, 1} ^*S*^ represents the distribution of wildtype (0) and alternative (1) alleles observed in the *S* samples. The conditional probability measure of the observations *y* given a realisation *x* is

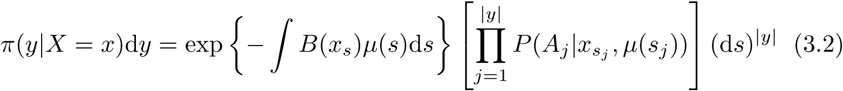

where *P* (*A*|*x*_*s*_, *µ*) is the probability of observing the allelic pattern *A* given a genealogy *x*_*s*_ and a mutation rate *µ* per nucleotide and per generation; this probably is calculated using Felsenstein’s peeling algorithm [19].

### 3.2. Choice of waypoints

To compute the minimum distance for waypoints, we need to compute the variance of the mutation rate before conditioning on data. For a complex model this is most straightforwardly estimated by simulation. Alternatively, assuming a panmictic population with constant diploid effective population size *N*_*e*_, the variance of the total coalescent branch length in a sample of *n* individuals is 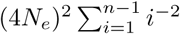 [58]. The variance of mutation rates *σ*^*2*^ is obtained by multiplying this by *µ*^2^, since the rate of mutations on the coalescent tree is *µ* times the total branch length. Rewriting this in terms of the heterozygosity *θ* = 4*N*_*e*_*µ*, and approximating the sum with 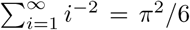 gives *σ*^2^ = *θ*^2^*π*^2^*/*6, and a minimum waypoint distance of 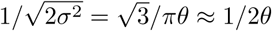.

### 3.3. The lookahead likelihood

The lookahead particle filter requires a tractable approximate likelihood of future data given a current genealogy. To achieve this we make a number of simplifications. First, most data are ignored, except for a digest of singletons and doubletons that are informative of the topology and branch lengths near the tips of the genealogy. This digest consists of the distance *s*_*i*_ to the nearest future singleton for each haploid sequence, and the *≤ n/*2 mutually consistent cherries (i.e. two-tipped nodes that can coexist in the same tree) *c*_*k*_ = (*a*_*k*_, *b*_*k*_), together with positions 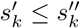 where their first and last supporting doubleton were observed. Under some simplifying assumptions we derive an approximation of the likelihood of the current genealogy given these data; see Section 5.5 for details.

### 3.4. Parameter inference

From (3.1) and Section 2.5, the parameters 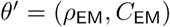 that maximize the expected log of 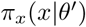, where the expectation is taken over *x* ∼ *π*(*x y, θ*)d*x* as approximated by algorithm 3, is

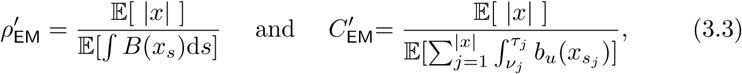

where *θ* = (*ρ, C*) is the vector of current parameter estimates. Note that 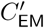 in (3.3) is constant in time. In practice we maximize (3.1) with respect to piecewise constant functions 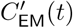, which yields

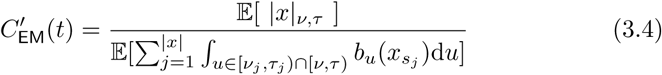

for *t* ∈ [*v, τ*), where |*x*| _*v,τ*_ denotes the number of coalescent events in *x* that occur in the epoch [*v, τ*). Similarly, a Variational Bayes inference procedure uses

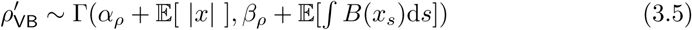

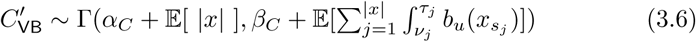

where expectations are taken over *x* ∼ *π*(*x*|*y, θ*)*p*(*θ*)d*θ*, where *p*(*θ*) is the posterior parameter distribution (3.5-3.6) of the previous iteration, and *α*_*ρ*_, *β*_*ρ*_, *α*_*C*_, *β*_*C*_ parameterize the prior distributions *ρ* ∼ Γ(*α*_*ρ*_, *β*_*ρ*_) and *C* ∼ Γ(*α*_*C*_, *β*_*C*_).

To estimate the expectations appearing in (3.3-3.6) we do not use full realisations *x* since resampling causes early parts of *x* to become degenerate due to “coalescences” of the particle’s sampling history along the sequence, resulting in high variance. Using the most recently sampled events is also problematic as these have not been shaped by many observations and mostly follow the prior *π*_*x*_(*θ*), resulting in highly biased estimates. Smoothing techniques such as two-filter smoothing [5] are difficult to apply here since finite-time transition probabilities are intractable. For discrete-time models fixed-lag smoothing is often used [45]. For our model the optimal lag depends on the epoch, as the age of tree nodes strongly influence their correlation distance. For each epoch we determine the correlation distance empirically, and for the lag we use this distance multiplied by a factor *α*; we obtain good results with *α* = 1.

### 3.5. Simulation study

We implemented the model and algorithm above in a C++ program `SMCSMC` (Sequential Monte Carlo for the Sequentially Markovian Coalescent) and assessed it on simulated and real data.

To investigate the effect of the lookahead particle filter, we simulated four 50 megabase (Mb) diploid genomes under a constant population-size model (*N*_*e*_ = 10, 000, *µ* = 2.5 *×* 10^*−*8^ / year, *ρ* = 10^*−*8^ / year, generation time *g* = 30 years). We inferred population sizes *N*_*e*_ = 1*/*(2*C*) through time as a piecewise constant function across 9 epochs (with boundaries at 400, 800, 1200, 2*k*, 4*k*, 8*k*, 20*k*, 40*k* and 60*k* generations) using particle filters Alg. 2 and Alg. 3, as well as a recombination rate, which was taken to be constant across time. Recombination rates were inferred accurately (data not shown) and throughout we focus on the accuracy of *N*_*e*_ through time. Observations are often available as unphased genotypes, and we assessed both algorithms using phased and unphased data, using the same simulations for both. Experiments were run for 15 EM iterations and repeated 15 times (Fig. 1a).

**Fig 1.**
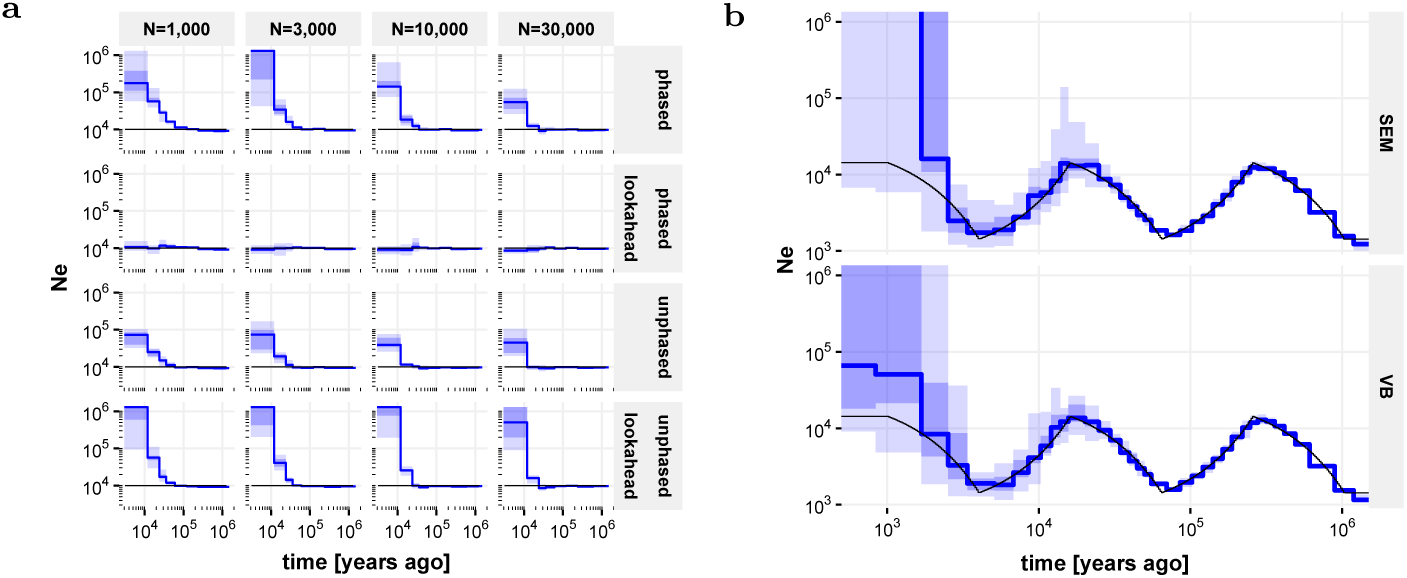
True population sizes (black), and median inferred population sizes across 15 independent runs (blue); shaded areas denote quartiles and full extent. **a** Impact of lookahead, phasing and number of particles on the bias in population size estimates for recent epochs, for data simulated under a constant population size model. **b** Inference in the “zigzag” model on phased data using lookahead and 30, 000 particles, comparing inference using stochastic Expectation Maximization (SEM) and Variational Bayes (VB).

On phased data (Fig. 1**a**, top rows), *N*_*e*_ values inferred without lookahead show a strong positive bias in recent epochs, corresponding to a negative bias in the inferred coalescence rate. Increasing the number of particles reduces this bias somewhat. By contrast, the lookahead filter shows no discernable bias on these data, even for as little as 1, 000 particles. On unphased data (Fig. 1**a**, bottom rows), the default particle filter continues to work reasonably well; in fact the bias appears somewhat reduced compared to phased data analyses, presumably because integrating over the phase makes the likelihood surface smoother, reducing particle degeneracy. By contrast, the lookahead particle filter shows an increased bias on these data compared to the default implementation. This is presumably because of the reliance of the lookahead likelihood on the distance to the next singleton; this statistic is much less informative for unphased data, making the lookahead procedure less effective, and even counterproductive for early epochs. Recombination rate inferences (assumed constant in time) were accurate in all experiments (data not shown).

We next investigated the impact of using Variational Bayes instead of stochastic EM, using the lookahead filter on phased data. We simulated four 2 gigabase (Gb) diploid genomes using human-like evolutionary parameters (*µ* = 1.25 *×* 10^*−*8^, *ρ* = 3.5 *×* 10^*−*8^, *g* = 29, *N*_*e*_(0) = 14312) under a “zigzag” model similar to that used in Schiffels and Durbin [51] and Terhorst, Kamm and Song [60], and inferred *N*_*e*_ across 37 approximately exponentially spaced epochs; see Section 5.6 for details. Both approaches give accurate *N*_*e*_ inferences from 2, 000 years up to 1 million years ago (Mya); other experiments indicate that population sizes can be inferred up to 10 Mya (data not shown; but see Fig. 2b). Inferences in the most recent epochs remain biased, which however is mitigated considerably by the Variational Bayes approach compared to SEM (Fig. 1b).

**Fig 2.**
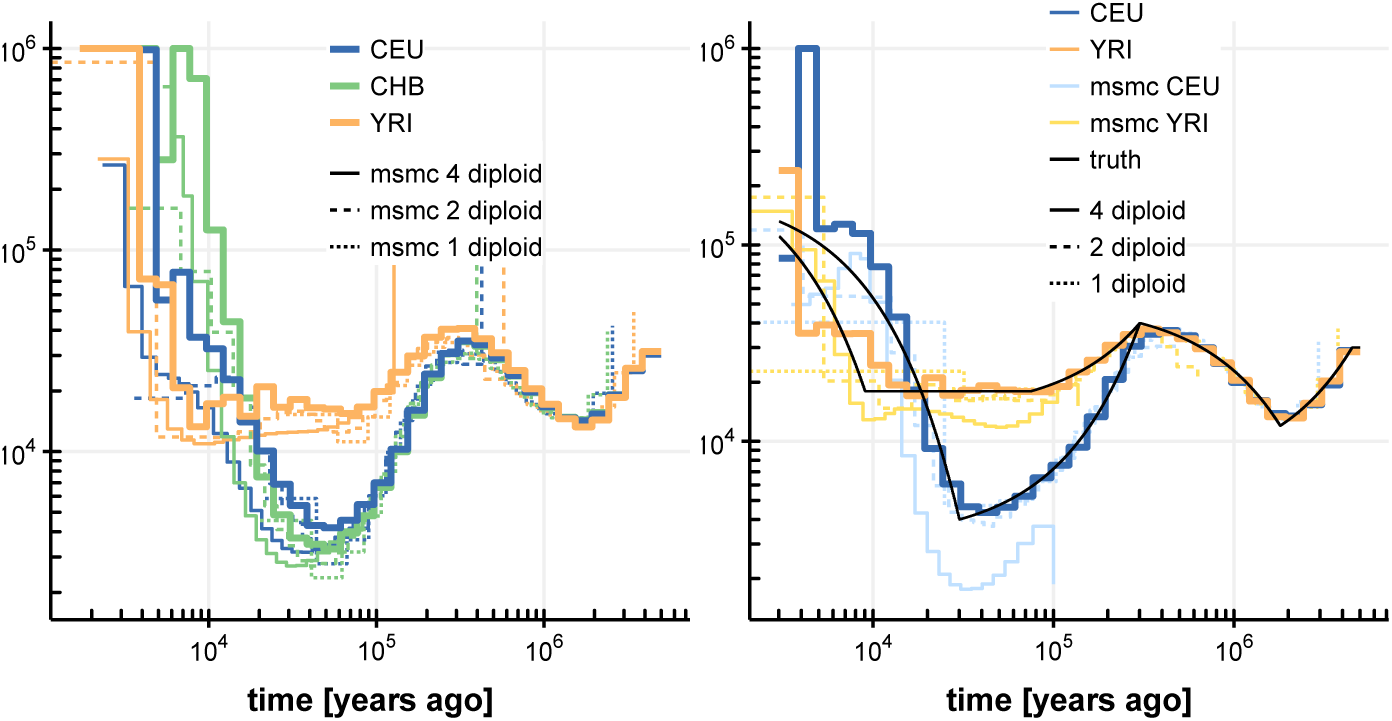
Population size inferences by `SMCSMC` on four diploid samples. Left, three human populations (CEU, CHB, YRI), together with inferences from `msmc` using 1, 2 and 4 diploid samples. Right, simulated populations resembling CEU and YRI population histories. All inferences (`SMCSMC`, `msmc`) were run for 20 iterations.

### 3.6. Inference on human subpopulations

We applied `SMCSMC` to three sets of four phased diploid samples of Central European (CEU), Han Chinese (CHB) and Yoruban (YRI) ancestry from the 1000 Genomes project. For comparison we also ran `msmc` [51] on the same data, and on subsets of 2 and 1 diploid samples. Inferences show good agreement where `msmc` has power. Since the inferences show some variation particularly in more recent epochs, we simulated data under a demographic model closely resembling CEU and YRI ancestry as inferred by multiple methods (see Section 5.6), and inferred population sizes using `SMCSMC` and `msmc` as before. This confirmed the accuracy of `SMCSMC` inferences from about 5,000 to 5 million years ago, while inferences in more recent epochs show more variability.

## 4. Discussion

Motivated by the problem of recovering a population’s demographic history from the genomes of a sample of its individuals [52], we have introduced a continuous-sequence approximation of the CwR model, and developed a particle filter algorithm for continuous-time Markov jump processes with a continuous phase space, by restricting the doubly-continuous process to a suitably chosen set of “waypoints”, and applying a standard particle filter to the resulting discrete-time continuous-state process. It however proved very challenging to obtain reliable parameter inferences for our intended application using this approach. To overcome this challenge we have extended the standard particle filter algorithm in two ways. First, we have generalized the Auxiliary Particle Filter of Pitt and Shephard [47] from a discrete-time one-step-lookahead algorithm to a continuous-time unbounded-lookahead method. This helped to address a challenging feature of the CwR model, namely that recent demographic events induce”sticky” model states with very long forgetting times. With an appropriate lookahead likelihood function (and phased polymorphism data), we showed that the unbounded-lookahead algorithm mitigates the bias that is otherwise observed in the inferred parameters associated with these recent demographic events. Some bias however remained, particularly for very early epochs. We reduced this remaining bias by a Variational Bayes alternative to stochastic expectation maximization (SEM), which explicitly models part of the uncertainty in the inferred parameters, and avoid zero rate estimates which are fixed points for the SEM procedure. In combination, the algorithm allowed us to infer demographic parameters from up to four diploid genomes across many epochs, without making model approximations beyond passing to the continuous-locus limit.

On three sets of four diploid genomes, from individuals of central European, Han Chinese and Yoruban (Nigeria) ancestry respectively, we obtain inferences of effective population size over epochs ranging from 5,000 years to 5 million years ago. These inferences agree well with those made with other methods [35, 53, 51, 57, 60], and show higher precision across a wider range of epochs than was previously achievable by a single method. Despite the improvements from the unbounded-lookahead particle filter and the Variational Bayes inference procedure, the proposed method still struggles in very recent epochs (more recent than a few thousand years ago), and haplotype-based methods [e.g., 29] remain more suitable in this regime. In addition, methods focusing on recent demography benefit from the larger number of recent evolutionary event present in larger samples of individuals, and the proposed model will not scale well to such data, unless model approximations such as those proposed in Terhorst, Kamm and Song [60] are used.

A key advantage of particle filters is that they are fundamentally simulation-based. This allowed us to perform inference under the full CwR model without having to resort to model approximations that characterizes most other approaches. The same approach will make it possible to analyze complex demographic models, as long as forward simulation (along the sequential variable) is tractable. The proposed particle filter is based on the sequential coalescent simulator SCRM [56], which already implements complex models of demography that include migration, population splits and mergers, and admixture events. Although not the focus of this paper, it should therefore be straightforward to infer the associated model parameters, including directional migration rates. In addition, several aspects of the standard CwR model are known to be unrealistic. For instance, gene conversions and doublet mutations are common [27, 62], and background selection profoundly shapes the heterozygosity in the human genome [41]. These features are absent from current models aimed at inferring demography, but impact patterns of heterozygosity and may well bias inferences of demography if not included in the model. As long as it is possible to include such features into a coalescent simulator, a particle filter can model such effects, reducing the biases otherwise expected in other parameter due to model misspecification. Because a particle filter produces an estimate of the data likelihood, any improved model fit resulting from adding any of these features can in principle be quantified, if these likelihoods can be estimated with sufficiently small variance. A further advantage of a particle filter is that it provides a sample from the posterior distribution of ancestral recombination graphs (ARGs). This allows estimating the age of mutations and recombinations, and explicit identification of sequence tracks with particular evolutionary histories, for instance tracts arising from admixture by a secondary population. In contrast to MCMC-based approaches [49], a particle filter can provide only one self-consistent sample of an ARG per run. However, for marginal statistics such as the expected age of a mutation or the expected number of recombinations in a sequence segment, a particle filter can provide weighted samples from the posterior in a single run.

The algorithm presented here scales in practice to about 4 diploid genomes, but requires increasingly large numbers of particles as larger numbers of genomes are analyzed jointly. This is because the space of possible tree topologies increases exponentially with the number of genomes observed, while the number of informative mutations grows much more slowly, resulting in increasing uncertainty in the topology given observed mutations. This uncertainty is further compounded by uncertainty in branch lengths.

Nevertheless, the many effectively independent genealogies underlying even a single genome provide considerable information about past demographic events [35], and a joint analysis of even modest numbers of genomes under demographic models involving migration and admixture events enable more complex demographic scenarios to be investigated. Our results show that particle filters are a viable approach to demographic inference from whole-genome sequences, and the ability to handle complex model without having to resort to approximations opens possibilities for further model improvements, hopefully leading to more insight in our species’ recent demographic history.

### 5. Appendix

#### 5.1. Conditional distributions and the Markov property

Here we outline how to define a conditional distribution *π*(*·*|𝒢) given a distribution *π* on *𝒳* and a conditioning subset *𝒢 ⊂ 𝒳* of measure 0. Suppose *𝒢* _*τ*_ is a family of subsets of *𝒳* so that *∪*_*τ*_ *𝒢* _*τ*_ =. A particular subset *𝒢* _*τ*_ for a fixed *τ* plays the role of the conditioning event *B* in the standard definition *P* (*A*|*B*) = *P* (*A ∩ B*)*/P* (*B*). It can be shown that, under some conditions, there exists an essentially unique family of measures 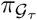 and a measure *µ* so that 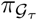 is concentrated on 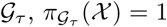 for all *τ*, and 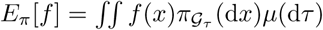 for well-behaved functions *f* [8], making it possible to define the conditional expectation as 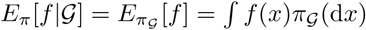. Using this, the Markov property of *π* can be expressed in terms of conditional expectations:

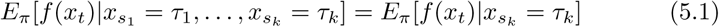

for loci *s*_1_ *< s*_2_ *<* … *< s*_*k*_ *< t* and any well-behaved function *f*.

#### 5.2. Proof of algorithm 3

The algorithm is proved by induction on *j*. For *j* = 0 the loop invariant is trivially true, while for *j* = *K* it implies the output condition. Suppose the loop invariant is true for some *j*. If *ESS < N/*2, assume w.l.o.g. that 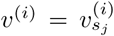 are normalized, let *i*_*k*_ be the index of the *k*th new particle, 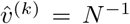 and 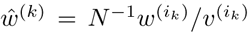 be its weights, and write 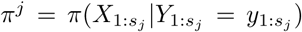, 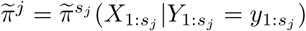 then

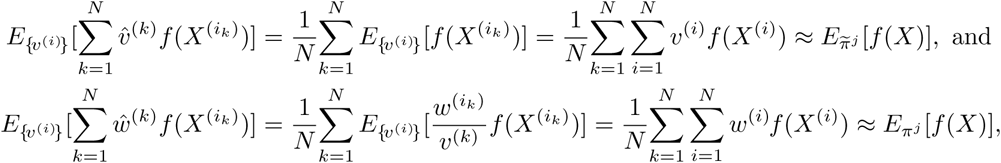

so that the loop invariant continues to hold after the optional resampling step.

After sampling 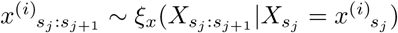, the particles 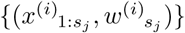 approximate 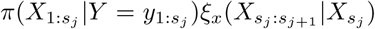. To make this distribution absolutely continuous w.r.t. 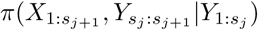, multiply it with the constant measure 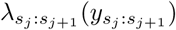; any measure will do as long as it has a density w.r.t. 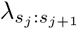 and is independent of *X*. Taking the Radon-Nikodym derivative of these two distributions gives

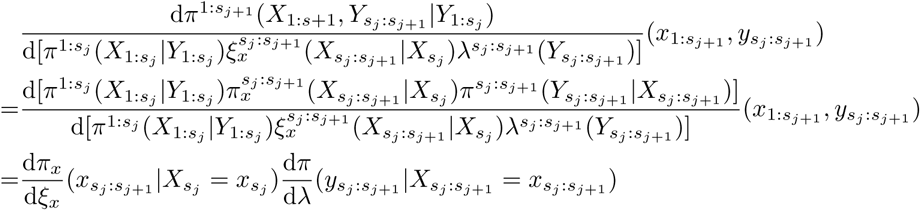

This shows that 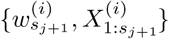 form particles approximating 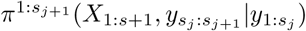 and since 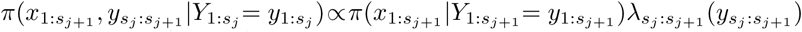 they also approximate 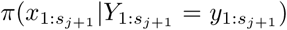. The argument showing that 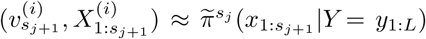 is analogous. This proves the loop invariant for *j* + 1, and the algorithm.

#### 5.3. The sequential coalescent with recombination process

To make formula (3.1) work, if *s* is a recombination point, *x*_*s*_ is the genealogy just left of the recombination point and includes the infinite branch from the root, so that *b*_*u*_(*x*_*s*_) = 1 for *u* above the root.

The distribution *π*_*x*_(*x*) has a density with respect to the Lebesgue measure (d*s*)^|*x*|^(d*u*)^2|*x*|^, because each of the |*x*| recombination events is associated with a sequence locus, a recombination time, and a coalescent time.

The measure (3.1) describes the CwR process exactly as long as *x* encodes both the local genealogy and the non-local branches used by the SCRM algorithm. In practice the SCRM algorithm prunes some of these branches, and we use (3.1) on the pruned *x*.

Note that we take the view that the realisation *x* encodes not only the sequence of genealogies *x*_*s*_ but also the number of recombinations |*x*| (some of which may not change the tree), their loci 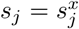, and the recombination and coalescence times 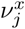 and 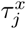. This information is also kept in the implementation of the algorithm, and is used to calculate the sufficient statistics required for inference of the coalescence and recombination rates.

#### 5.4. Variational Bayes for Markov Jump processes

We consider hidden Markov models where the latent variable follows a Markov jump process over *x* ∈ 𝒳, that with respect to a suitable measure d*x*d*y* admits a probability density of the form

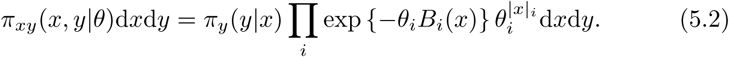

Here, |*x*| _*i*_ is the event count for events of type *i* in realisation *x*, and *B*_*i*_(*x*) is the total opportunity for events of that type in *x*. For example, in our case

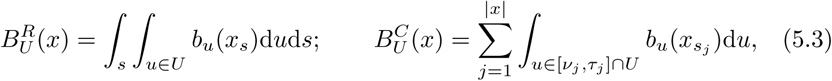

and 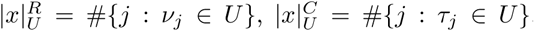, for recombinations and coalescence opportunities and counts occurring in an epoch *U ⊂*[0, ∝).

A Variational Bayes approach approximates the true joint posterior density *π*(*x, θ*|*y*) ∝ *π*_*xy*_(*x, y*|*θ*)*π*_*θ*_(*θ*), where *π*_*θ*_ is a prior on the parameters, with a probability density *φ*(*x, θ*) that is easier to work with (here the constant of proportionality implied by “∝ “hides a constant density *λ*(*y*)). Following Hinton and van Camp [31] and Mackay [37], we choose to constrain *φ* by requiring it to factorize as *φ*(*x, θ*) = *φ*_*x*_(*x*)*φ*_*θ*_(*θ*), and we choose to optimize it by minimizing the Kullback-Leibler divergence *KL*(*φ π*), also referred to as the variational free energy [20],

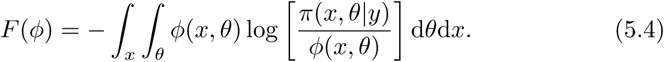

To optimize *φ*_*θ*_(*θ*) we write *F* (*φ*) as a function of *φ*_*θ*_ with *φ*_*x*_ fixed, as

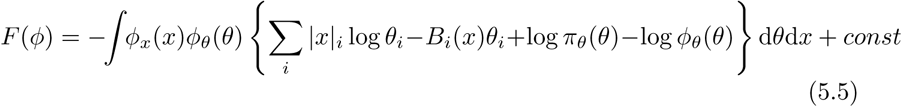

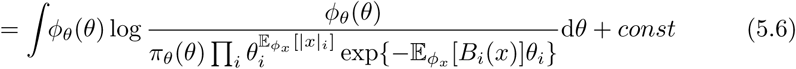

This is minimized by setting log *φ*_*θ*_(*θ*) equal to the denominator. We can still choose the prior *π*_*θ*_(*θ*); a product of Gamma distributions 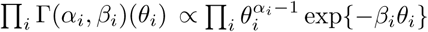 is suitable as it is conjugate to the factors appearing in the denominator. The result is that

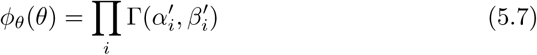

with 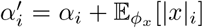 and 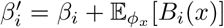. Next, to optimize *φ*_*x*_(*x*) we write *F* (*φ*) as a function of *φ*_*x*_ with *φ*_*θ*_ fixed,

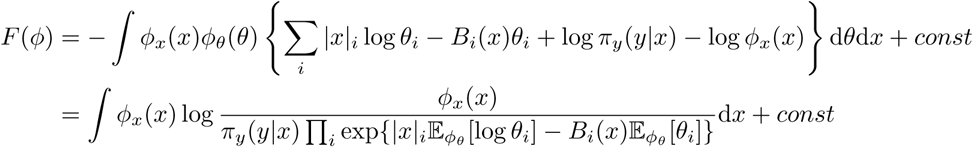

Define 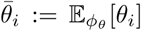 and 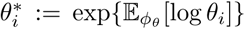, then using properties of the Gamma distribution we get 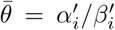 and 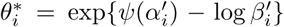 where *Ψ* is the digamma function. Again, *F* (*φ*) is minimized if the numerator and denominator are equal, which happens for

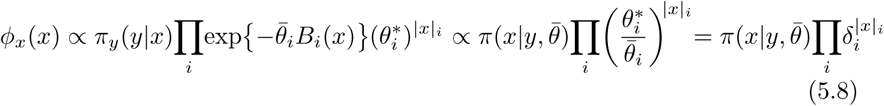

where 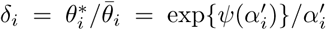. As given, the algorithms in this paper sample from a distribution of the form 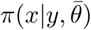, but they can easily be modified to sample from *φ*_*x*_(*x*) instead by including an additional factor *δ*_*i*_ in a particle’s weight for every event of type *i* that occurs.

#### 5.5. Approximate lookahead likelihood

Let *s*_*i*_ denote the distance to the nearest future singleton in each sequence, and let *c*_*k*_ = (*a*_*k*_, *b*_*k*_) be *≤n/*2 mutually consistent cherries with positions 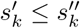 of their first and last supporting doubleton. To simplify notation we assume that the current position is 0 (Fig. 3a).

**Fig 3.**
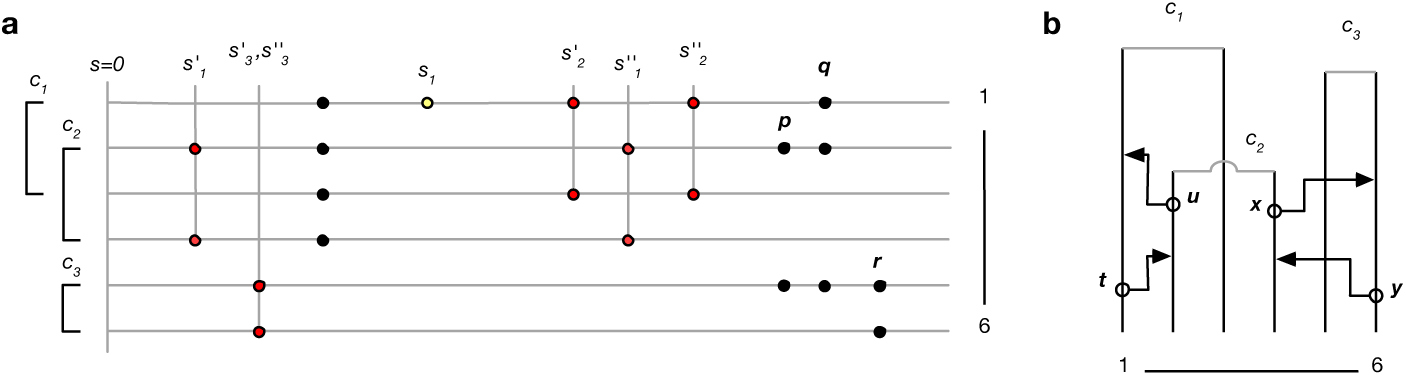
**a**. Example of data digest. Lines represent genomes of 6 lineages, circles observed genetic variants. Of the data shown, one singleton (yellow) and five doubletons (red) contribute to the digest. Cherry c_3_ is supported by a single doubleton; r does not contribute because the mutation patterns p and q are incompatible with c_3_. Similarly, p does not contribute because it is incompatible with c_2_ and c_3_. **b**. Partial genealogy (unbroken lines) over 6 lineages. Open circles and arrowheads represent potential recombination and coalescence events that would change the terminal branch length for lineage 1 (t,u), and remove cherry 3 (x,y).

Note that recombinations result in a change of a terminal branch length (TBL) if either the recombination occurred in the branch itself and the new lineage does not coalesce back into it, or the recombination occurred outside the branch and the new lineage coalesces into it (Fig. 3b). To compute the likelihood that the first singleton in lineage *i* occurs at position *s*_*i*_, we assume that all TBLs are equal to *l*_*i*_, and that coalescences occur before *l*_*i*_. Then, the total rate of events that change the TBL *i* is

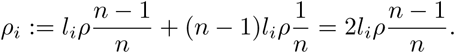

Define *µ*_*i*_ := *µl*_*i*_ to be the total mutation rate on branch *i*, and assume that when a TBL changes, it reverts deterministically to some length 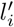. If a terminal branch with length *l*_*i*_ changes at *u* to 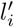, which happens with probability 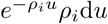, the likelihood that the first singleton occurs at distance *s*_*i*_ is 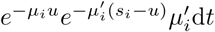, where 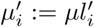. Conversely, if that branch does not change along [0, *s*_*i*_), which happens with probability 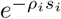, the likelihood is 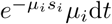. Combining these possibilities and marginalizing over *u ∈* [0, *s*_*i*_) gives

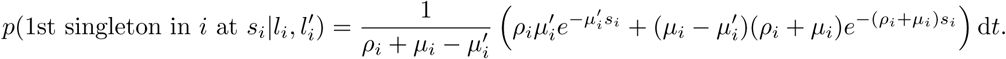

In the case that no singleton is observed up until *s*_*i*_ but data was missing thereafter, the same probability densities apply except for the factors *µ*_*i*_ and 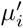 in the likelihood, so that

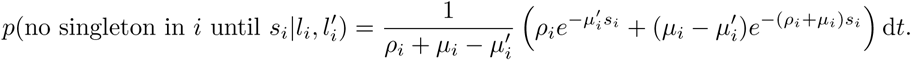

We account for the uncertainty in 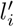 by marginalizing over the empirical distribution of TBLs for sequence *i*.

To approximate the likelihood of the doubleton data, note that a cherry *c* = (*a, b*) at height *l* changes if a recombination occurs in either branch *a* or *b* and the new lineage coalesces out, or a recombination occurs outside of *a* and *b* and coalesces into either (Fig. 3b). Again assuming that all TBLs are *l* and coalescences occur before *l*, the total rate of change is 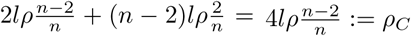. When a cherry changes, we assume that the new cherry is drawn from the equilibrium distribution. To calculate the probablity of observing *c* = (*a, b*) at equilibrium, assume that a tree supports 1 *≤ k ≤ n/*2 cherries. The branches of *c* are among the 2*k* branches subtended by the tree’s *k* cherries with probability 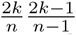 and *a* is paired with *b* with probability 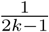. Since *k* has mean *n/*3 if *n ≥* 3 [39], the probability of observing (*a, b*) at equilibrium is 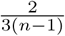. We approximate the likelihood of a doubleton by 0 if the *c* is not in the tree, and by 1 if it is. Then, the likelihood of observing *c*_*k*_ = (*a*_*k*_, *b*_*k*_) at the last known position 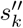 conditional on the tree currently containing *c*_*k*_ is

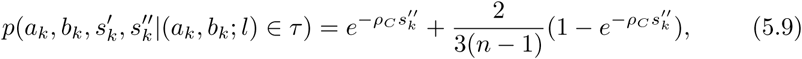

where (*a*_*k*_, *b*_*k*_; *l*) ∈ *τ* expresses that *τ* contains cherry *c*_*k*_ = (*a*_*k*_, *b*_*k*_) at height *l*. Now suppose *c*_*k*_ */∈ τ* and let 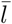 be the average TBL in *τ*. Under similar assumptions, cherries are created at a rate 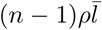 and assuming that new cherries are drawn from the equilibrium distribution, the likelihood of observing *c*_*k*_ at the first known position 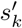 is

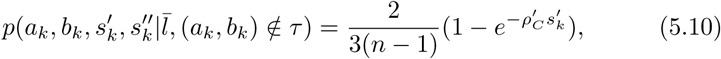

where 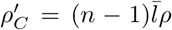 is the effective rate of recombinations that potentially result in the creation of *c*_*k*_. Note that (5.9-5.10) are likelihoods for *τ* supporting *c*_*k*_ at the given position, rather than for a doubleton mutation actually occurring.

These likelihoods show good performance, but result in some negative bias in inferred population size for recent epochs. We traced this to the lack of correlation between *l*_*i*_ and 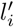, requiring a single very recent coalescence to explain a long segment devoid of singletons, rather than allowing for the possibility of several correlated coalescences each in slightly earlier epochs. To model correlations, we averaged the likelihood above over *ρ*′ = *ρ* and *ρ*′ = *ρ*/2 each weighted with probability 1*/*2. This effectively removed the negative bias.

To deal with missing data, we reduce *µ* proportionally to the missing segment length and the number of lineages missing. For unphased mutation data, singletons and doubletons can still be extracted, and are greedily assigned to compatible lineages. The likelihoods are also similarly calculated, by greedily assigning cherries to observed doubletons. Unphased singletons can result from mutations on either of the individual’s alleles; the likelihood term uses the sum of the two branch lengths for that individual to calculate the expected rate of unphased singletons.

#### 5.6. Implementation details

While *x*_1:*s*_ refers to the entire sequence of genealogies along the sequence segment 1 : *s*, storing this sequence would require too much memory. Instead we only store the most recent genealogy *x*_*s*_ (including non-local branches where appropriate), which is sufficient to simulate subsequent genealogies using the SCRM algorithm. To implement epoch-dependent lags when harvesting sufficient statistics, we do store records of the events (recombinations, coalescences and migrations) that changed *x* along the sequence, as well as the associated opportunities, for each particle and each epoch; this implicitly stores the full ARG. To avoid making copies of potentially many event records when particles are duplicated at resampling, these are stored in a linked list, and are shared by duplicated particles where appropriate, forming a tree structure. Records are removed dynamically after contributing to the summary statistics, and when particles fail to be resampled, ensuring that memory usage is bounded.

The likelihood calculations involve many relatively slow evaluations of the exponential function, often for small exponents. We use the continued-fraction approximation 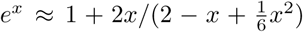 for |*x*| *<* 0.03, with relative error bounded by 10^*−*10^ [36].

Table 1 shows the commands to generate the data for the three simulation experiments. Epoch boundaries for *N*_*e*_ inference in generations for the zigzag experiment were defined by taking interval boundaries − 14312 log(1 −*i/*256)*/*2, *i* = 0, …, 255, merging intervals according to the pattern 4 *\**1 + 7 *\**2 + 8 *\**5 + 7 *\**13 + 1 *\** 15 + 8 *\**11 + 1 *\**3 (37 epochs; see [35]). For the real data experiments, epochs boundaries for the 32 epochs were logarithmically spaced from 133 to 133016 generations ago, using generation time *g* = 29 years, without merging intervals (command line option `-P 133 133016 31*1`).

**Table 1.** Commands to generate simulation data

| Experiment | Command |
| --- | --- |
| zigzag | `scrm 8 1 -N0 14312 -t 1431200 -r 400736 2000000000 -eN 0 1 -eG 0.000582262 1318.18 -eG 0.00232905 -329.546 -eG 0.00931619 82.3865 -eG 0.0372648 -20.5966 -eG 0.149059 5.14916 -eN 0.596236 0.1 -seed 1 -T -L -p 10 -l 300000` |
| CEU | `scrm 8 1 -N0 14312 -t 1789000 -r 500920 2500000000 -eN 0 10.4807 -eG 0.00120468 214.8965 -eG 0.0180702 -14.15827 -eG 0.180702 1.33255 -eG 1.084212 -0.563414 -eN 2.71053 2.096143 -seed 1 -T -L -p 10 -l 300000` |
| YRI | `scrm 8 1 -N0 14312 -t 1789000 -r 500920 2500000000 -eN 0 10.4807 -eG 0.00120468 502.8635 -eG 0.00542106 0 -eG 0.0451755 -5.89189 -eG 0.180702 1.33255 -eG 1.084212 -0.563414 -eN 2.71053 2.096143 -seed 1 -T -L -p 10 -l 300000` |

